# A Longitudinal Analysis of Function Annotations of the Human Proteome Reveals Consistently High Biases

**DOI:** 10.1101/2024.10.18.619148

**Authors:** An Phan, Parnal Joshi, Claus Kadelka, Iddo Friedberg

## Abstract

The resources required to study gene function are limited, especially when considering the number of genes in the human genome and the complexity of their function. Therefore, genes are prioritized for experimental studies based on many different considerations, including, but not limited to, perceived biomedical importance, such as disease-associated genes, or the understanding of biological processes, such as cell signaling pathways. At the same time, most genes are not studied or are under-characterized, which hampers our understanding of their function and potential effects on human health and wellness. Understanding function annotation disparity is a necessary first step toward understanding how much functional knowledge is gained from the human genome, and toward guidelines for better targeting future studies of the genes in the human genome effectively. Here, we present a comprehensive longitudinal analysis of the human proteome utilizing data analysis tools from economics and information theory. Specifically, we view the human proteome as a population of proteins within a knowledge economy: we treat the quantified knowledge of the protein’s function as the analog of wealth and examine the distribution of information in a population of proteins in the proteome in the same manner distribution of wealth is studied in societies. Our results show a highly skewed distribution of information about human proteins over the last decade, in which the inequality in the annotations given to the proteins remains high. Additionally, we examine the correlation between the knowledge about protein function as captured in databases and the interest in proteins as reflected by mentions in the scientific literature. We show a large gap between knowledge and interest and dissect the factors leading to this gap. In conclusion, our study shows that research efforts should be redirected to less studied proteins to mitigate the disparity among human proteins both in databases and literature.

## 1 Introduction

One of the most important and challenging endeavors in the genomic era is to provide a deep functional understanding of the amount of available genetic sequence data [1–3]. At the same time, allocating resources to study all available genes and gene products equally is inefficient and practically infeasible. Therefore, prioritizing which genes to study is an essential goal of genome analysis and annotation [4], and there are several criteria that researchers may use to prioritize genes for functional studies. These criteria can be phenotypical, such as prioritizing genetic disease-related genes, antimicrobial resistance genes, or quantitative trait loci for crop enhancement [5–9], or phylogenetic, such as focusing on highly conserved gene families for evolutionary genomics [10, 11]. Additionally, genes may be prioritized due to their involvement in key biochemical pathways, such as kinases, or regulatory roles in gene expression networks, such as transcription factors [12–14].

Previous studies have shown a sustained bias in which genes are chosen for functional studies. Edwards *et al* (2011) noted that “researchers’ favorite kinases have remained the same for decades, with a few exceptions” [12]. The study made apparent a research preference towards a few well-studied kinases, and hypothesized several reasons for this phenomenon. First, scientists may feel more skilled and therefore are more comfortable working within their area of expertise, rather than opening new avenues of research. Second, most funding and peer-review systems are averse to unsupported areas of research, viewing them as inopportune risks, and unwilling to support research on unstudied proteins. Third, academic promotion culture and results-driven demands incentivize quick outcomes, favoring familiar research methods and well-established research infrastructure [12]. A subsequent study identified that research into gene function is “primarily guided by a handful of generic characteristics of genes facilitated by experimentation during the 1980s and the 1990s, rather than physiological importance of individual genes or their relevance to human disease” [15]. Due to these favorable characteristics, certain genes were studied more intensively in the pre-genomic era and have attracted substantial attention ever since [16]. With the introduction of high-throughput sequencing, many novel genes were discovered in the last two decades, providing a more comprehensive view of the human genome. However, the historical bias persists and these newly identified genes receive less attention and follow-up studies than previously favored genes, despite preliminary evidence of their importance [17]. Further, some genes were found to be abandoned in a “leaky” pipeline between the findings of –omics experiments and the writing of manuscripts. Specifically, less studied genes are less likely to be highlighted in the title or abstract despite being significant –omics hits because authors are not incentivized to report them [18]. Taken together, despite many advances in the discovery of new genes, subsequent and necessary experiments to characterize the functions these genes remain lacking.

The present literature discussing human genes and proteins reflects an imbalance in functional studies. Previous studies have characterized this bias in science literature, concluding that a small group of proteins, well-characterized and studied for decades, continue to attract an overwhelming fraction of life science research articles [16, 19–21]. Biases in functional studies of genes are also reflected in the inequality of their Gene Ontology annotations, in which certain genes lack specific annotations and result in less effective subsequent research on less annotated genes [12,20]. The Gene Ontology (GO) is a standardized semi-hierarchical vocabulary consisting of defined terms and relationships that annotate the molecular function, biological process, and cellular component aspect of a gene or protein [22]. Functions of genes and proteins are described with GO annotations. Any inequality among GO annotations of different genes hints at a disparity in our understanding of gene function. This inequality in GO annotations has been shown to be growing over time in Human and model organisms [23]. Importantly, understanding gene function and annotation inequality in model organisms is no less important than in Human. This is because much of our understanding of human gene function and systems is inferred from model organisms, and a bias in the latter will contribute to undetected biases in the former [24, 25].

The study and the annotation of genes and proteins have a sustained bias with disproportionate focus on well-studied proteins linked to specific diseases, leaving many proteins with potential therapeutic relevance understudied and under-annotated [26]. This inequality creates gaps in our understanding and potential treatments for complex conditions, impacting overall knowledge about genes and proteins involved in human health [19,20]. Addressing these disparities is crucial for advancing our comprehensive biomedical discoveries and improving health outcomes [26, 27]. Recognizing the problem of understudied genes and proteins, and several initiatives have been taken to mitigate the inequality in the study of gene and protein function [20, 28–30]. In 2017, the “Illuminating the Druggable Genome” project was launched to discover target genes that express proteins able to bind drug-like molecules, where the project focused on three protein families: protein kinases, ion channels, and G-protein-coupled receptors [26, 31]. More recently, the “Find My Understudied Genes” software was developed, which encouraged researchers to report lesser-known genes in their publications and design experiments around these understudied genes [21]. These efforts aimed to redistribute research focus towards those genes with poorly characterized or unknown function, and subsequently reduce the inequality in protein function studies. Therefore, examining the inequality of protein functions over time becomes essential to offer deeper and up-to-date insights into the biases in the study of proteins and the annotation of protein function. In our work, we examine the inequality of protein studies and protein functions overtime to assess the impact of previous initiatives, measure the current level of inequality, and to provide recommendations for further prioritization of genes for functional research.

Previous studies have explored the inequality in literature mentions as well as GO annotations among human proteins; however, no direct correlation between these two measures has been established. In this study, we examine the correlation between the publication mentions and GO annotations of human proteins. In doing so, we note that accurately quantifying the information represented in the GO annotations for each protein is crucial. Traditional methods often simply count the number of GO terms assigned to the proteins [23, 32]. However, this approach does not account for the varying informativeness of GO terms. For instance, a “child” GO term such as `acyltransferase activity`, which describes the catalysis of the transfer of an acyl group, is more informative than the “parent” GO term `transferase activity`. Building on previous publications that primarily quantified the knowledge of protein functions by counting their GO annotations, we apply an information-theory-based approach that quantifies the information represented by each GO term. Our approach utilizes the informativeness of GO annotations to more precisely quantify the knowledge of protein functions, resulting in a more accurate quantification of knowledge about protein function [32–35].

Here, we take a novel approach to reviewing inequality in protein function by analogizing the knowledge about protein function to personal wealth. Specifically, we characterize *knowledge* of a protein’s function as a quantifiable asset that grows “richer” with informative annotations achieved from experiments. Given this measure of knowledge wealth for a single protein, we can extrapolate it to encompass the “knowledge wealth” for a whole population of proteins, in this case, the human proteome [23]. Viewing the knowledge of protein function via a quantitative economic lens allows us to perform an analysis analogous to the study of wealth inequality using classic economic tools [36]. The motivation of this study is to gain a better insight into the changing inequality in the knowledge of proteins over time, acknowledging where the “wealth” predominantly resides and how it should be better distributed.

The adapted information-theoretic and economic tools allow us to perform a comprehensive longitudinal analysis of annotation inequality in the human proteome. Our findings underscore the ongoing biases in the function annotation of human proteins. Additionally, we use economic metrics to measure the inequality of knowledge from these proteins over a decade, revealing that the inequality of protein annotations has remained consistently high from 2013 to 2022. Addressing these disparities is crucial for advancing our biomedical knowledge and improving health outcomes. Furthermore, our study explores the correlation between the overall attention or interest level a protein receives and the extent of our knowledge of its functions. We show that there is a weak correlation between the number of publication mentions of proteins and the quantified knowledge from their GO annotations. We also show that there is still a large gap between literature-stored knowledge and database knowledge. Our findings highlight the critical need to refocus efforts on less studied proteins to mitigate the annotation inequality effectively and to develop more ways to capture functional data from papers in a computationally amenable way.

## 2 Methods

The Gene Ontology (GO) knowledgebase is a large and comprehensive compilation of information on the functions of genes and gene products. GO terms are organized in a hierarchical structure, with more general terms at higher levels and more specific terms at lower levels. This allows for the description of gene products at varying levels of specificity, which can be quantified. GO can be seen as a means to capture functional knowledge of proteins in a human and computationally readable manner. Gene products (such as proteins or RNAs) are annotated with GO terms, which indicate their roles in biological processes, molecular functions, and cellular components (which are the three aspects of the GO). The annotations are based on experimental evidence or computational predictions.

First, we quantified the available knowledge of each protein as captured in the GO. Here, *knowledge* means the amount of data we have about any protein’s function: the different types of functions, and the level of specificity, or precision of a certain function, all of which we derived from the GO. We therefore used the following metrics to capture GO-based knowledge: term count, unique term count, and information content, which will be described in the following subsections. Taken together, these metrics capture the knowledge about each protein that is represented in the GO.

Besides knowledge, we also wanted to capture the interest in a protein. Here, *interest* means how much the protein is studied, which may not necessarily correlate with how much we know about a protein. To measure interest, we used the normalized number of scholarly publications mentioning the protein. Additionally, we examined the Spearman correlation between the knowledge and interest metrics.

Finally, we looked at the inequality among the proteins with respect to the knowledge and interest metrics. To do so, we used the Gini coefficient, which is an economic measure of dispersion commonly used to represent income inequality in a population (e.g., a country) [36]. We adopted the Gini coefficient to represent the inequality in a protein population, in this case, the Human proteome [23]. The analog of income or wealth here would be the knowledge of or, conversely, the interest in a protein. Following this summary, we describe the different method parts in more detail.

### 2.1 Knowledge metrics

A key aspect of this study is understanding how much “knowledge wealth” each protein has accrued over time and comparing this wealth between different proteins. One dimension of wealth that can be quantified is the functional knowledge available for any given protein. We used the GO knowledgebase as the initial source of annotation to quantify knowledge for each human protein in the UniProt Knowledgebase (UniProtKB-SwissProt) [22, 37, 38]. GO annotation (GOA) files are released on an approximately monthly basis and stored in the GO Data Archive https://release.geneontology.org/index.html. We used annotations with experimental evidence codes (EXP, IDA, IPI, IMP, IGI, IEP, HTP, HDA, HMP, HGI, HEP). We further categorized the annotations based on their source articles (using the “Reference” column in the GOA files). In this study, we used GOA files from December of each year between 2013 and 2022. For GO structures, we used the .obo files from the GO Data Archive at the same time points. To quantify the amount of knowledge we have of a protein’s function, we used the following metrics.

#### 2.1.1 Term count and unique term count

Term count is the number of most specific GO terms assigned to each protein. A GO term is considered the most specific for a particular protein if no other GO term assigned to the protein is its descendant [35]. If a protein is annotated with the same GO term more than once with different source data (e.g., different articles) and/or a different associating partner in the “With” column of a GOA file (e.g., one protein binds many partner proteins), each annotation is counted towards term count. Therefore, although duplicated annotations might be considered redundant information, they are not discarded in the term count metric. The rationale for keeping duplicated GO terms lies in the observation that if the annotations are recorded in different entries, it may reflect (1) the attention in that particular function as it comes from different sources with different evidence codes, or (2) a variation of the function (e.g., a different binding partner). Let *𝓛_x_* denote the collection of most specific GO terms of protein *x*, term count is therefore *|𝓛_x_|*. In our analysis, we also used unique term count, a conventional metric for quantifying GO terms [23, 32], which is the number of unique GO terms in *𝓛_x_*.

#### 2.1.2 Information content

GO terms vary in specificity, or in how informative they are. E.g., `kinase activity` is less informative than `protein tyrosine kinase activity`. Since term count and unique term count do not capture how informative a GO term is, we used an information-theory based approach to correctly quantify the informativeness of a GO term. To do so, we adapted the method of using GO term frequency as a basis for the information content of a GO term, as described in [34,39]. Specifically, each GO term *i* has an information content (IC) based on the frequency *P_i_* of that term in the human proteome. Adapted from Claude Shannon’s presentation of information content [33], since information content is linked to term frequency, a less frequent term has higher surprisal (or information content) if a protein is annotated with it [40].

Each protein *x* is annotated to all GO terms in *𝓛_x_* (in GOA files). For each protein, we built its annotation directed acyclic graph (DAG), denoted *T_x_*, by collecting all GO terms in *𝓛_x_*, and including ancestral terms by propagating each term up to the root via the `is_a` and `part_of` edge types, where each node or edge in the resulting DAG appears once. The information content of a protein *x*, in bits, can be computed from *T_x_*associated with the protein as follows.

- Following Clark *et al*, we model the relationships between GO terms in the ontology using a Bayesian network and thus assume each GO term is independent of its ancestors, given its parents [39]. Under this assumption, the probability of any protein being annotated with a sub-graph *T* of the ontology, denoted as Pr(*T*), can be calculated by multiplying the conditional probabilities of each node *v* (or GO term *v*) in *T* given its parent nodes 𝓟(*v*) as follows:

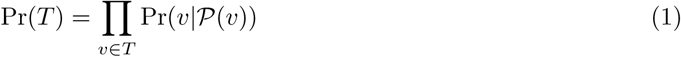 Here, Pr(*v*|𝓟(*v*)) represents the conditional probability of a protein being annotated with GO term *v*, given that the protein is already annotated with the parent GO terms in 𝓟(*v*).
- The information content of a sub-graph *T* of the ontology, denoted *i*(*T*), is calculated as:

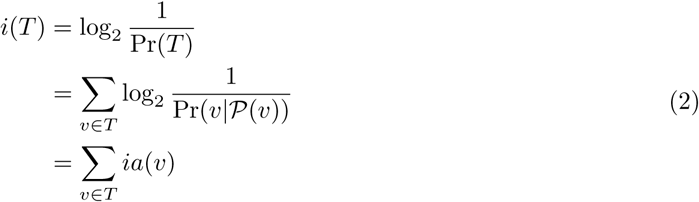

where 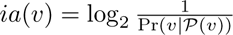 represents the *information accretion* of term *v*. The information accretion of term *v* can be thought of as the additional information gained about the sub-graph *T* by including the term *v*, beyond the information already provided by the parent terms 𝓟(*v*). *i*(*T*) is thus the sum over all the information accretions *ia*(*v*) for all terms *v* in *T* . See [39] for further details.
- The information content of any protein *x*, annotated with the sub-graph *T_x_*, is thus the information content of *T_x_*, or *i*(*T_x_*). It is worth noting that each protein has three annotation DAGs, one for each aspect of the GO; therefore, each protein has three information content values, one for each GO aspect.

The information content of GO terms and proteins changes with each Uniprot-GOA release, so the information content values are different between releases due to the change in (1) the structure of the Gene Ontology DAG (e.g., addition/obsoletion of a GO term, or addition/deletion of a relationship between two GO terms) and (2) the change in the frequency of annotations in the corpus [41]. To standardize our analysis across time, we used the Gene Ontology DAG and GO term frequencies from December 2022 as standards for calculating the information content of proteins. First, we collected most specific GO annotations for each protein each year from 2013 to 2022. Second, we constructed the annotation DAG for each protein using the Gene Ontology in 2022. Third, we calculated the information accretion of GO terms using the 2022 GO structure and GO annotations of proteins. Finally, we computed the information content of each protein by summing up the information accretion values (obtained from the third step) of every term in the protein’s annotation DAG (constructed in the second step) following Eq (2). These steps enable a comparison of the information content of proteins across years. Additionally, we noticed that certain proteins lost annotations from one year to the next, which may be due to (1) the annotated term becoming obsolete and being removed from the GO, or (2) a biocurator correcting a past error, resulting in the protein no longer being annotated to that term. Since this is a technical issue that does not advance our understanding of knowledge gain over time, we set all negative information content gain values to zero. In other words, when calculating the gain in information content of a protein from 2013 to 2022, the proteins cannot lose information.

### 2.2 Interest metrics

In addition to quantifying the knowledge of proteins based on GO annotations, we examined another aspect that reflects how extensively human proteins are studied in the scientific literature. We used an *interest metric* to quantify the attention a protein receives within the scientific community. To quantify the interest in a protein, we computed its *total fractional count*, as described by Sinha *et al* [19]. Briefly, the fractional count *f_xd_* of protein *x* in publication *d* is the ratio between how many times the protein *x* is mentioned in the publication and the total number of mentions of all proteins in that publication. The fractional count was computed for each protein in each publication. The total fractional count *f_x_* of protein *x* is the sum of all its fractional counts across all considered publications. That is,

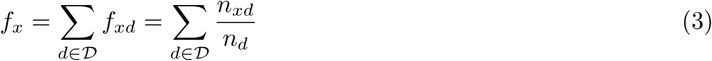

where *𝓓* is the document set, *n_xd_* is the number of times protein or gene *x* is mentioned in document *d*, and *n_d_* is the total number of mentions of any protein or gene in document *d*. Eq (3) describes the total amount of literature published about a protein *x* across all publications in *𝓓*, consisting mostly of available full-text articles or abstracts from PubMed. Sinha *et al* introduced the term *full publication equivalent* as a unit for the total fractional count metric. We quantified the interest for any given protein using the number of FPEs of that protein [19]. We obtained the number of FPEs of all human proteins from 2013 to 2022 from the Pharos/TCRD database, which is part of the Knowledge Management Center project (https://download.jensenlab.org/KMC/) . Note that the number of FPEs of proteins in 2013 is accumulated from publications before 2013. The gain in the number of FPEs per protein was calculated by subtracting the number of FPEs per protein in 2022 from that in 2013. It should be noted that some proteins that are mentioned in publications not because they are subjects of research, but because they are mostly or exclusively used as assaying or biomedical diagnostic tools, for example, albumin, or insulin.. We therefore excluded 109 proteins that are biomarkers, identified in MarkerDB [42] from our analysis, as the majority of the publications mentioning these proteins are clinical reports that do not contribute new annotations to protein function. Additionally, we removed CD4 and TNF due to their exceedingly high number of FPEs (greater than 70,000 FPEs), also primarily attributable to their widespread use in clinical or diagnostic assays [43, 44]. While it was not feasible to eliminate all proteins used as biomarkers or in clinical assays, these exclusions aimed to minimize the impact of these types of outliers in the FPE data, ensuring a more accurate analysis.

### 2.3 Measuring inequality

Next, we wanted to examine the inequality of the knowledge and interest among proteins in the human proteome. To do so, we used the *Gini Coefficient*. The Gini coefficient is typically used to measure the inequality in the income distribution in populations of countries or other geopolitical units [36, 45]. We adopted the Gini coefficient to measure the inequality in the distribution of annotations among a set of proteins. To do so, we used the GO term count, unique GO term count, and information content as an analog of wealth and the human proteome as the “population”. For each of the three knowledge metrics, the intuition is the same: a protein is “wealthier” in knowledge if the knowledge metric is higher. We then calculated each year’s Gini coefficient for the population of proteins to determine how the inequality in function annotation of human proteins changes over time. In this calculation, we include the proteins with a term count that is greater than zero at a particular time point, i.e., proteins without any annotations in any given GO aspect at that time point are not included in the population.

### 2.4 Correlation between knowledge and interest

Having introduced knowledge and interest as different aspects of “protein wealth”, we wanted to discover the relationship between knowledge and interest over time. We computed the pairwise correlations between four metrics: the number of FPEs of all proteins in 2013, the information content of all proteins in 2013, their gain in number of FPEs from 2013 to 2022, and their gain in information content from 2013 to 2022. The rationale behind these correlation tests is to predict the gain in the number of FPEs and the gain in information content after ten years using the knowledge and interest metrics in the past, in this case, starting in 2013. All analyses used Spearman-ranked correlation.

Additionally, we used the DISEASES database for disease-gene associations [46] to determine if proteins related to certain diseases have a stronger link between interest and knowledge. We focused on diseases that have at least 20 proteins associated with them. We then performed Spearman correlation tests between the number of FPEs of proteins within each disease and their gain in information content (separately in Molecular function and in Biological process aspects of the GO) from 2013 to 2022. We controlled the family-wise error rate at 0.05 using the Bonferroni multi hypothesis correction [47, 48].

### 2.5 Time delay between publication and annotation

We also examined the time delay between publication and curation of protein functions to better understand the gap between community research interest and database-stored knowledge. We retrieved the publication year of all PubMed articles that were used to annotate human proteins experimentally, which we called the publication timestamp for each annotation. Next, we extracted the curation timestamp from the “Date” field in GOA files for each annotation. Note that we only included the earliest instance of the GO annotation for the same protein (e.g., different protein binding partners) for repeated annotations to the same protein as we favor primary annotation results [49]. In other words, only unique protein-GO term pairs are included in the analysis of the delay in curation. Additionally, we removed annotations sourced from the IntAct database, which consist of `protein binding` and `identical protein binding` terms. The reason for removing IntAct annotations is that the curation timestamp is updated with every GOA file and does not reflect the timestamp in which the GO term was first assigned to a protein. Finally, the delay in curation for a given annotation is the number of years between the publication timestamp and the curation timestamp.

## 3 Results

### 3.1 Our knowledge about protein function is gradually increasing, and specific proteins show bursts of information gain

We used GO term count, GO unique term count, and information content to capture different facets of knowledge about protein function. We also considered the type of publication from which a GO annotation was derived. Specifically, we differentiated between articles describing high-throughput experiments (articles annotate more than 100 proteins per article) and articles describing low-throughput experiments (articles annotate no more than 100 proteins per article). This distinction is made because previous publications have shown that while large-scale experiments often annotate many proteins, they annotate each protein with fewer unique GO terms, which are also less informative than the terms that medium- and low-throughput experiments annotate the proteins [2,50–52]. Moreover, the annotations generated from large-scale experiments usually require additional validation in smaller-scale experiments [53, 54].

First, we quantify the knowledge gained from protein annotations using information content. As depicted in Figure 1A, articles describing high-throughput experiments consistently yield less informative annotations for the proteins, which aligns with previous findings [50, 55]. The average information content of proteins studied in low-throughput experiments is much greater than those studied in high-throughput experiments.

**Figure 1:**
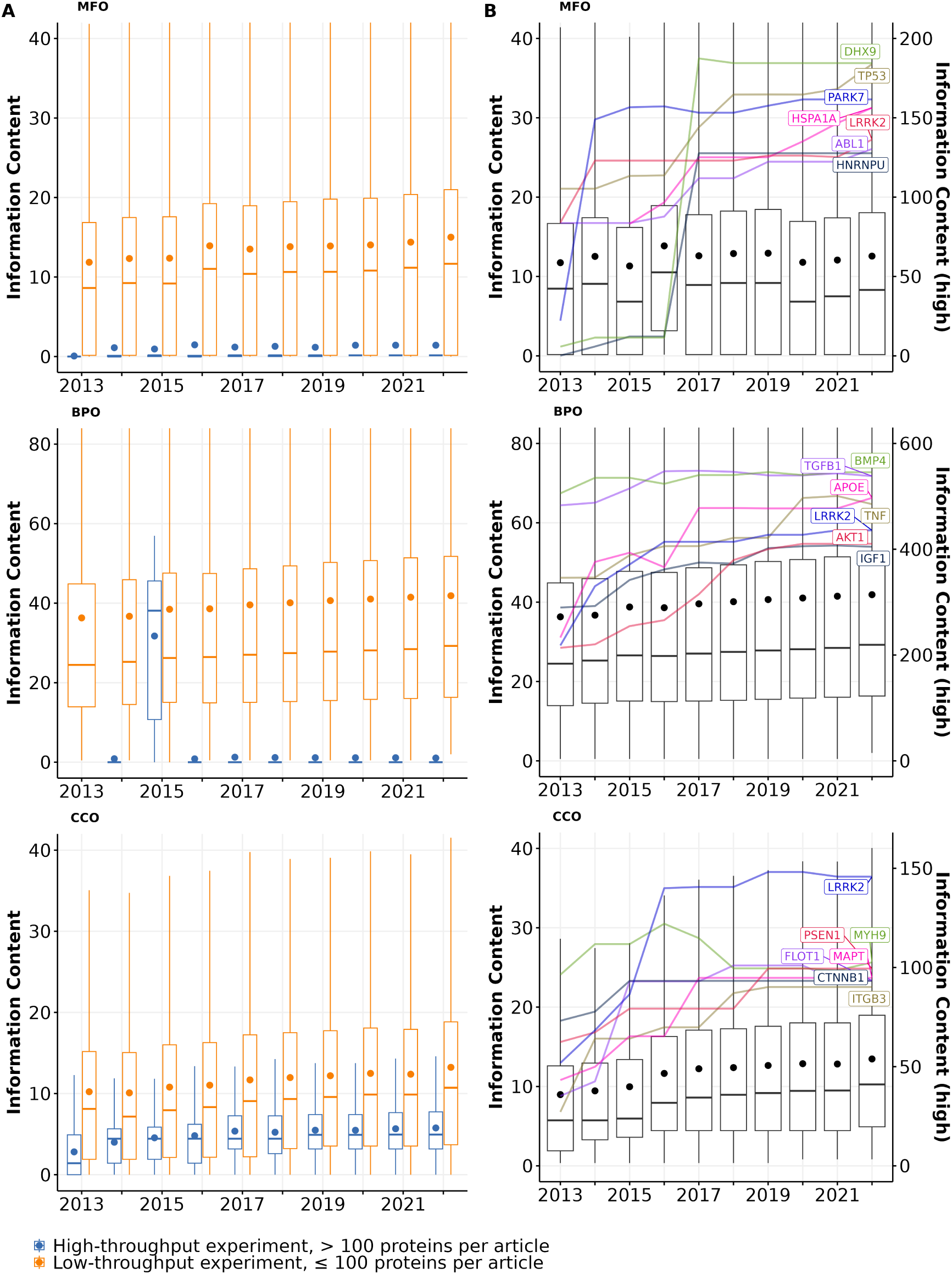
Information content of proteins studied experimentally from 2013 to 2022 in (A) articles describing high-throughput (> 100 proteins per article; blue boxes) vs. low-throughput experiments (*≤* 100 proteins per article; orange boxes) and (B) all articles. Each box shows the distribution of information content for all proteins with a given year and aspect: MFO (Molecular function), BPO (Biological process), and CCO (Cellular component). Each box extends across the interquartile range (IQR), vertical lines extend to the lowest data point (or highest data point) still within 1.5 IQR of the lower quartile (or the upper quartile), a horizontal line shows the median and a dot shows the mean value. Outliers are not displayed. In panel B, highlighted lines show the growth of information content for the most highly annotated proteins over time. Their information content values are described by the right y-axis (see Table S2 for gene and protein names).

For the proteins that received annotations from both article types in Molecular function (MFO) and Cellular component (CCO), the majority of their information content was contributed by articles describing low-throughput experiments. Notably, about 20% of proteins with experimental MFO or CCO annotations were exclusively annotated using high-throughput experiments, yet these annotations carried little information. In contrast, only a small number of Biological process (BPO) annotations that proteins received were derived from high-throughput experiments, and by the end of 2022, most of these annotations have been removed. Only three proteins (Q15661, Q14155, and P23946) have their experimental BPO annotations derived from articles describing high-throughput experiments.

The overall information content of human proteins is expected to increase over time as we discover more functions of these proteins, expand the Gene Ontology by adding more specific GO terms, and continuously curate available information [56, 57]. Figure 1B shows that the median information content of all proteins increased over time for BPO (from 24.5 to 29.2) and CCO (from 5.7 to 10.3), which was driven by the knowledge gained from both proteins annotated before 2013 and after 2013 (see Table S1). Overall, both the number of experimentally annotated proteins and the median information content of proteins in BPO and CCO aspects increased steadily, implying that we are accruing more knowledge about human proteins annotated before 2013 and adding annotations for newer proteins through experiments over time. Interestingly, the median information content of proteins in the MFO aspect fluctuated but did not increase. This is due to a large number of proteins that are newly annotated every year between 2013 and 2022 whose only annotation is `protein binding`. When proteins with `protein binding` as sole annotation are excluded, the median information content of proteins in the MFO aspect increased slightly from 13.6 to 14.8 (see Figure S1).

Figure 1B also highlights the best-annotated proteins with the highest information content by the end of 2022. Among the best-annotated proteins, we observe two distinct trends in information content growth within the MFO aspect: some proteins exhibited a striking information content growth rate over a short period, while others showed a steady, less dramatic increase. For instance, PARK7, a gene known for its protective role in Parkinson’s disease, showed a remarkable six-fold increase in experimental information content between 2013 and 2014 [5,6].The burst increase in information content of PARK7 is likely attributed to the UK Parkinson’s Disease Consoritum (UKPDC) Initiative which started in January 2014, resulting in both focused research efforts and annotation efforts [58]. Similarly, two well-annotated multi-functional proteins HNRNPU and DHX9 also show substantial growth between 2016 and 2017 with 10-fold and 16-fold increases in MFO information content, respectively [59, 60]. Other proteins, such as TP53, HSPA1A, APOE, AKT1, gained information content more gradually over time. Notably, some proteins like LRRK2 in MFO and BPO, as well as BMP4 and TGFB1 in BPO, expressed a growth followed by a plateau, suggesting a potential saturation of information that can be quantified from their GO annotations. The observed saturation of information observed with some proteins could be because (1) we did not gain more new knowledge about the function of these proteins, or (2) the knowledge we acquired about a protein’s function belongs to specific areas of biology that are not in the scope of the Gene Ontology (e.g., addition or modification of a protein’s functional groups through post-translational modifications, or disease associations) [41].

Next, we quantified the knowledge gained from protein annotations through unique term count and term count. We found that articles describing high-throughput experiments generally assign fewer unique MFO and CCO terms than those describing low-throughput experiments, whereas in the BPO aspect, most annotations were derived from articles describing low-throughput experiments (Figure S2) [50]. In contrast, when using the term count metric (i.e., the same GO term is counted as many times as they are recorded in GOA files), annotations from high-throughput experiments make up two-thirds of the annotations in MFO and one-third in CCO. The majority of annotations that are repeatedly assigned to proteins are mostly: `protein binding` and `RNA binding` in MFO aspect (each annotation specifies a binding partner), `nucleus`, `cytoplasm`, `cytosol` in CCO aspect (each annotation specifies a different source article). Although these terms have low information content, they describe information that is not captured in the information content metric, such as details about binding partners or different supporting articles. Notably, high-throughput experiments were responsible for a substantial fraction of MFO and CCO annotations for new proteins, i.e., proteins that did not have experimental annotations previously in these two aspects. From 2013 to 2022, articles describing high-throughput experiments exclusively annotated 43% of the new proteins with MFO annotations (2957 out of 6800 new proteins) and 34% of the new proteins with CCO annotations (1688 out of 5004 new proteins). Therefore, while articles of high-throughput experiments offer fewer unique and less informative annotations per protein, they play an important role in providing initial knowledge about the functions of proteins that have thus far not been studied in detail experimentally [53, 61].

In Table 1, we list the top 10 proteins with the highest information content or highest term count in each aspect.

**Table 1:**
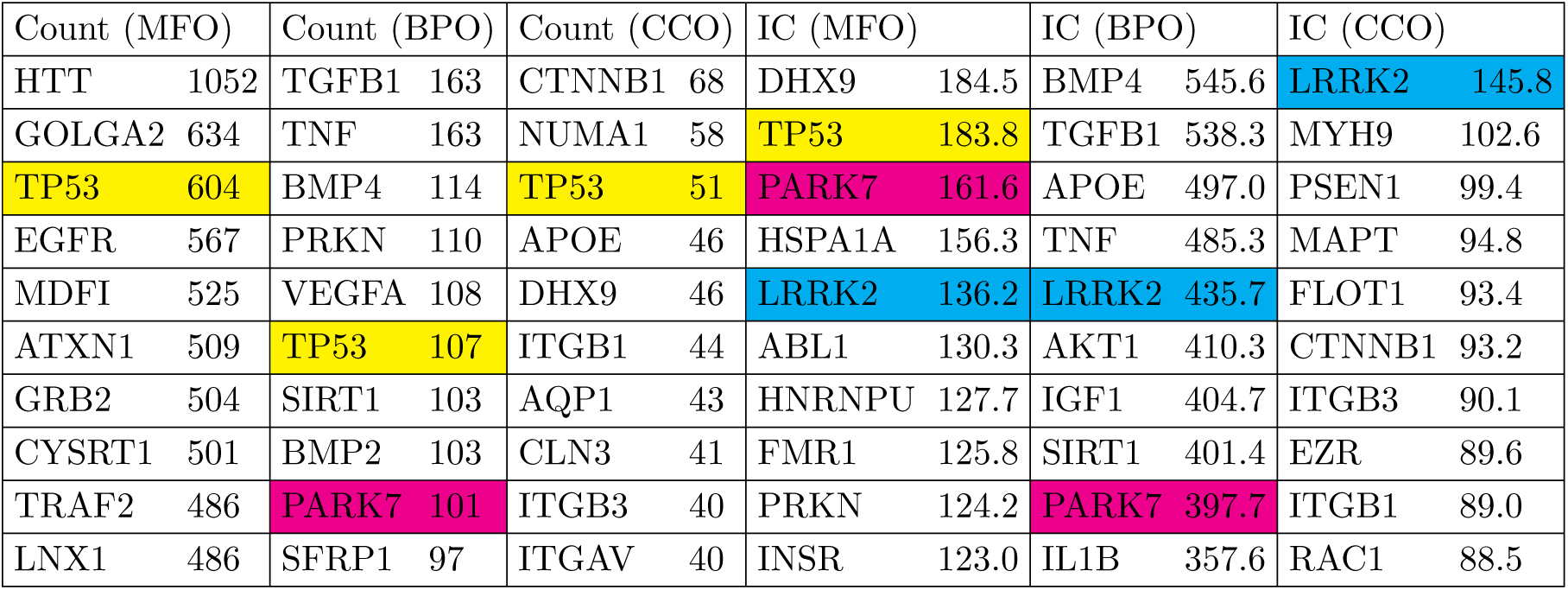
Top 10 proteins with highest term count or highest information content (IC) in each GO aspect (MFO, BPO, CCO). Proteins that appear at least three times in the table are highlighted. Data retrieved from the GOA files released in Dec 2022.

Table 1 shows the ten proteins with the highest term count and highest information content for each GO aspect. The gene HTT, coding for the protein Huntingtin, has the highest MFO term count of 1052 experimental annotations among all human proteins. However, the information content that this protein gained from 900 annotations from large-scale experiments is notably low at 5 bits, most of which being `protein binding` but with different binding partners. Meanwhile, low-throughput experiments provided Huntingtin with more than 330 bits of information via highly informative annotations. TP53 has a high term count for all three aspects, especially for MFO, with roughly 500 `protein binding` annotations and 35 other unique GO annotations. However, TP53 does not have one of the highest information content values in BPO and CCO. This highlights that relying solely on term count provides an insufficient quantification of the available knowledge about protein function.

In each aspect, we possess the most knowledge for a different protein, indicating that certain proteins are extensively annotated in a particular aspect than in others. For instance, DHX9 is known for its multi-functional role as ATP-dependent RNA helicase A, which interacts with both DNA and RNA and is involved in post-transcriptional regulation; these characterized functions are highly specific, which gives DHX9 the highest information content in MFO [62, 63]. Similarly, BMP4 — bone morphogenetic protein 4 — plays a crucial role in numerous developmental processes and BMP pathways [64]. LRRK2 — leucine-rich repeat serine/threonine-protein kinase 2 — not only is an enzyme involved in a broad range of processes (e.g., neuronal plasticity, innate immunity, autophagy), but is also found to reside in highly specific subcellular locations [65, 66].

Figure 1B also reveals that although the average information content of proteins increases gradually over time, the spread of this distribution did not change. Specifically, the information content distributions have consistently been right-skewed with large interquartile ranges. A few proteins are extreme outliers (i.e., their information content values are very high, see 1), while the information content of most proteins is low. These observations suggest a high inequality in the amount of information that proteins have. Furthermore, there is no shrinkage in the interquartile range from 2013 to 2022 for all three GO aspects, implying that the imbalance in function annotation of proteins persisted from 2013 to 2022.

### 3.2 Inequalities in annotations and publications remain high over time

To assess the imbalance in function annotation between highly annotated proteins and less annotated proteins, we calculated the Gini coefficients of the knowledge distribution of human proteins. The Gini coefficient is adopted here to quantify annotation inequalities, where higher Gini coefficients indicate a higher level of annotation inequality [36, 45]. Specifically, we treat term count, unique term count, and information content as “income” measures to evaluate the inequality in function annotation of human proteins over time. We calculated the Gini coefficient for each year between 2013 and 2022. All experimentally annotated proteins constitute the protein population (see Table S1).

Figure 2A reveals that all knowledge metrics derived from GO annotations of human proteins in every GO aspect show consistently high Gini coefficients over a ten-year span from 2013 to 2022. The Gini coefficients of every metric in every GO aspect range from 0.4 to 0.65, with small disparities across years, indicating substantial and persistent inequalities in the annotation of human proteins. In particular, the high Gini coefficients of 0.6-0.65 for term count in MFO imply that proteins receive varying number of annotations, primarily due to the frequent and repeated annotation of many proteins to the `protein binding` term. Notably, the consistently high Gini coefficients of 0.6 for the distribution of MFO information content suggest a large disparity in information content among these proteins. This is largely because 36% of proteins only have one MFO annotation of `protein binding`, which carries a low information content of 0.15 bits. When repeated GO terms are counted only once, as in the unique term count metric, the Gini coefficient drops to approximately 0.4. Although unique term count per protein was shown to correlate strongly with information content per protein (Figure S3, [32]), our analysis shows that the distribution of information content is more unequal than the distribution of unique term count in the MFO aspect. Due to the prevalence of low-information “protein-binding-only” proteins, it is most meaningful to use the information content metric to quantify the available knowledge about proteins in MFO aspect, instead of unique term count. Unlike the MFO aspect with high and increasing inequality in term count and information content, the Gini coefficients for the knowledge metrics in BPO and CCO aspects remained at lower values (ranging from 0.4 to 0.5) and mostly unchanged over time. There is also less disparity between the Gini coefficients calculated from term count, unique term count, and information content in these two aspects. Despite calls for more comprehensive function annotation, inequalities measured by Gini coefficients of the different metrics have not notably decreased between 2013 and 2022 [12, 15, 67, 68] (see Figure S4).

**Figure 2:**
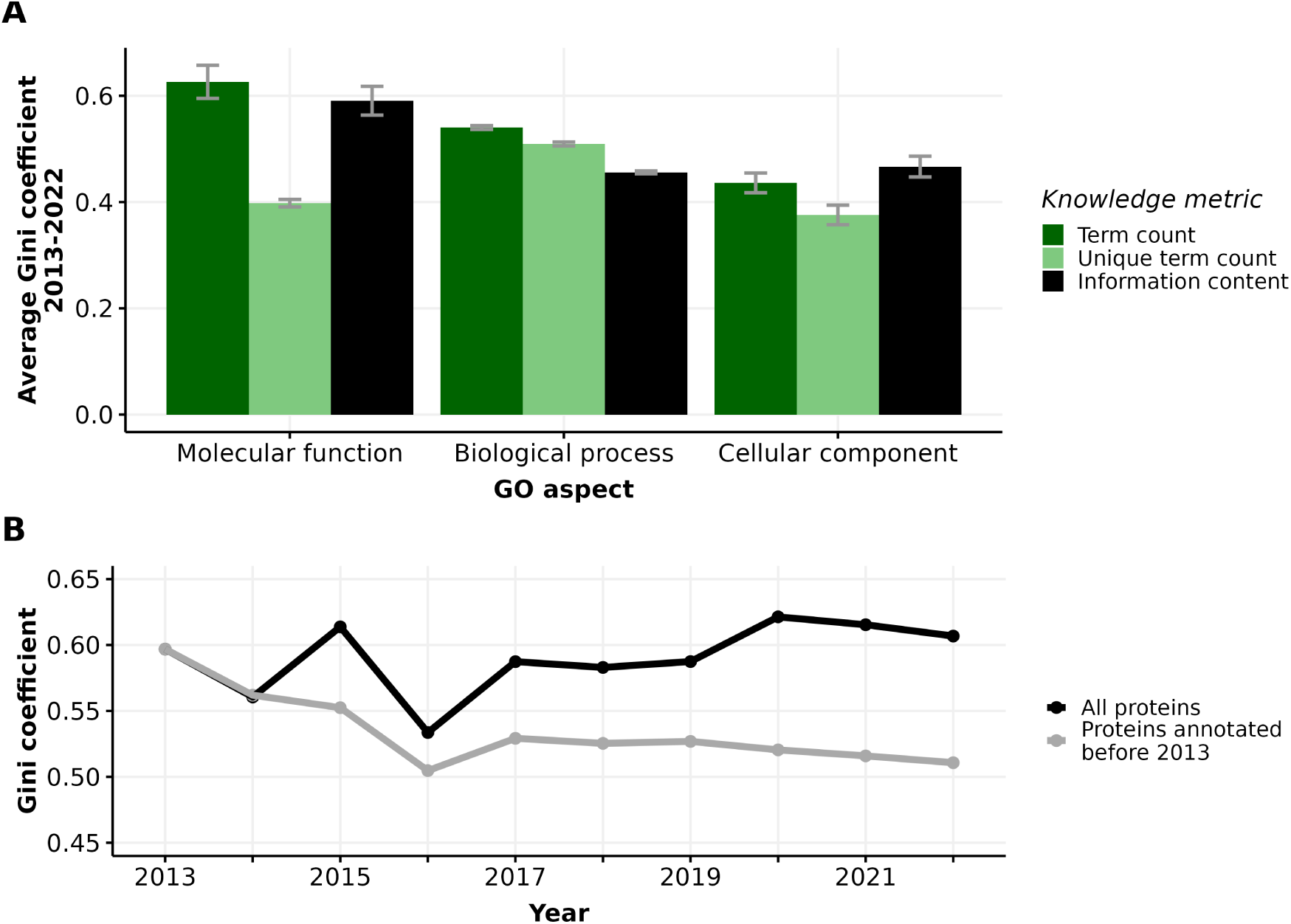
(A) Barplot displaying average Gini coefficient (0-1) from 2013 to 2022. Each bar indicates one of three knowledge metrics of human proteins: term count (dark green), unique term count (light green), and information content (black) in one of three GO aspects. Error bars extend one standard deviation above and below the average value. (B) Gini coefficient from 2013-2022 calculated from information content in Molecular function (MFO) aspect of two subsets of proteins: all proteins (black) and proteins first annotated before 2013 (grey).

The number of annotated proteins in each aspect increases significantly (Table S1) from 2013 to 2022, facilitating our comparison of all proteins versus proteins annotated in or before 2013. Considering the set of 7,831 proteins that were annotated experimentally in or before 2013, the Gini coefficient for the distribution of information content in MFO actually decreased from 0.59 to 0.51 from 2013 to 2022 (Figure 2B, Table S1). Unlike the fluctuating Gini coefficients for the MFO information content of all proteins, the decreasing trend of the protein subset annotated in or before 2013 could be driven by more even annotation efforts for this subset. As some of these proteins had already been annotated extensively and became well-annotated proteins, resources could focus on other proteins that were annotated in or before 2013 with less informative annotations. Specifically, we show that the distribution of information content in MFO of proteins annotated in or before 2013 is evidently less right-skewed than that of proteins annotated after 2013 (Figure S5), confirming that earlier-annotated proteins are more equally annotated, likely due to longer research periods into their function. Even though the Gini coefficients of MFO information content of all proteins fluctuated from 2013 to 2022, we show that the subset of earlier-annotated proteins experienced a decrease in inequality, indicating that we are accruing more knowledge about earlier-annotated proteins as well as discovering novel information about recently-annotated proteins at the same time. For the information content in BPO and CCO aspects, there is almost no difference in Gini coefficients between all proteins and the subset of earlier-annotated proteins (see Figure S4).

Parallel to our knowledge about protein function, research interest in human proteins is also highly unequal. The number of FPEs per human protein quantifies the interest each protein receives over time. The Gini coefficients of the number of FPEs are extremely high, at 0.87 in 2013 and have been decreasing to 0.85 in 2022 (Figure 3). Note that the number of proteins in this analysis is greater than the number of experimentally annotated proteins in the knowledge metric analysis. The inclusion of many proteins with low number of FPEs and without any experimental annotation (i.e., these proteins were excluded from the calculation of Gini coefficient for information content metric) may contribute to the substantially higher Gini coefficient in interest versus knowledge.

**Figure 3:**
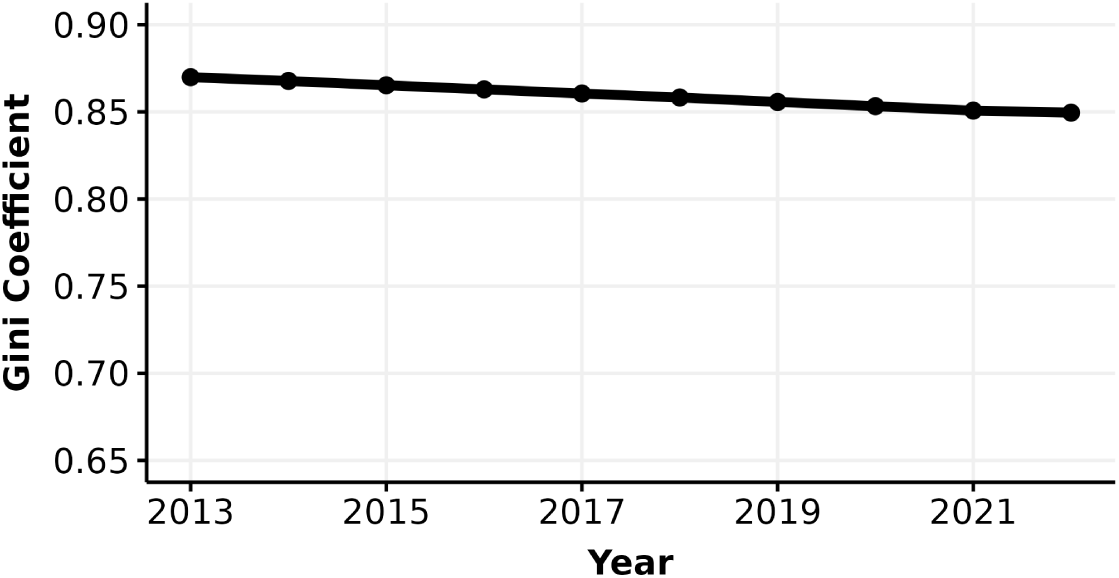
Gini coefficient (0-1) of the number of FPEs of human proteins.

We found that the median number of FPEs of human proteins has increased from 13.7 to 34.2 and the mean number of FPEs increased from 143.1 to 277.1 from 2013 to 2022, suggesting that the distribution of the number of FPEs is consistently right-skewed [19](see Figure S6). Furthermore, the number of FPEs in 2013 is strongly correlated with the gain in number of FPEs from 2013 to 2022 (*ρ*_Spearman_ = 0.94, *p* < 3 *×* 10^−16^, Figure S7). This strong correlation suggests that the number of FPEs of proteins in 2013 is highly predictive of the gain in number of FPEs after 10 years, and thus proteins with higher number of FPEs in 2013 gained more FPEs compared to proteins with lower number of FPEs.

Notably, we did not observe the same longitudinal correlation in the case of information content of human proteins. The Spearman coefficients between information content in 2013 and the gain in information content after 10 years were negative (*ρ*_Spearman_ = −0.24 for MFO and *ρ*_Spearman_ = −0.41 for BPO, with *p <* 3*×*10^−16^ for both tests). While the initial interest in proteins is correlated with the gain in interest, initial knowledge of proteins is not indicative of the gain in knowledge. This result suggests that although proteins that were mentioned more frequently in the past tend to accumulate more publications, we are not gaining a corresponding amount of information about their functions.

### 3.3 Research interest is not indicative of the gain in knowledge

Following the analysis on knowledge of and interest in proteins as different aspects of “protein wealth”, we discovered how correlated they are. We examined the correlation between interest, as represented by the number of FPEs, and knowledge, as represented by information content. Figure 4 shows a weak correlation between information content and the number of FPEs of proteins over a 10-year period. By highlighting proteins with the largest growth in information content over 10 years (similar to Figure 1), we notice that a few proteins (DHX9, PARK7, HNRNPU) gain a large amount of information content and are among the most annotated proteins, but they do not receive that much interest. We also highlighted four proteins with all-time high interest, reflected by high number of FPEs: ALB (albumin), CRP (C-reactive protein), INS (insulin), and IL6 (interleukin-6). Unlike proteins with highest growths in information content, these high interest proteins did not gain information content over time while their number of FPEs increases substantially. The reason is that ALB, CRP, INS, and IL6 are mostly used as a biomedical tool and being mentioned in a large amount of clinical reports, rather than being researched for function [49,69–72]. In line with this, insulin and interleukin-6 showed signs of saturated information content in the MFO aspect, while C-reactive protein and albumin reached saturation in the BPO aspect. The signs of saturation in information content of the high interest proteins suggested that these proteins were not annotated with new functions, which is consistent with their appearance in literature not as research subjects, but as an assay component. After removing the proteins that are primarily mentioned as assay components in publications (see Methods), we computed the Spearman correlation between the information content and the number of FPEs per protein for each year between 2013 and 2022. We found that the Spearman correlation coefficients between the number of FPEs per protein and the information content per protein are in the ranges of (0.39, 0.42), (0.31, 0.34), and (0.17, 0.24) in the aspect of MFO, BPO, and CCO, respectively. By the end of 2022, the correlation coefficients are quite low for all experimentally annotated proteins (*ρ*_Spearman_ = 0.42, 0.34, 0.17 for MFO, BPO, CCO, respectively, with *p <* 3 *×* 10^−16^ for all three tests).

**Figure 4:**
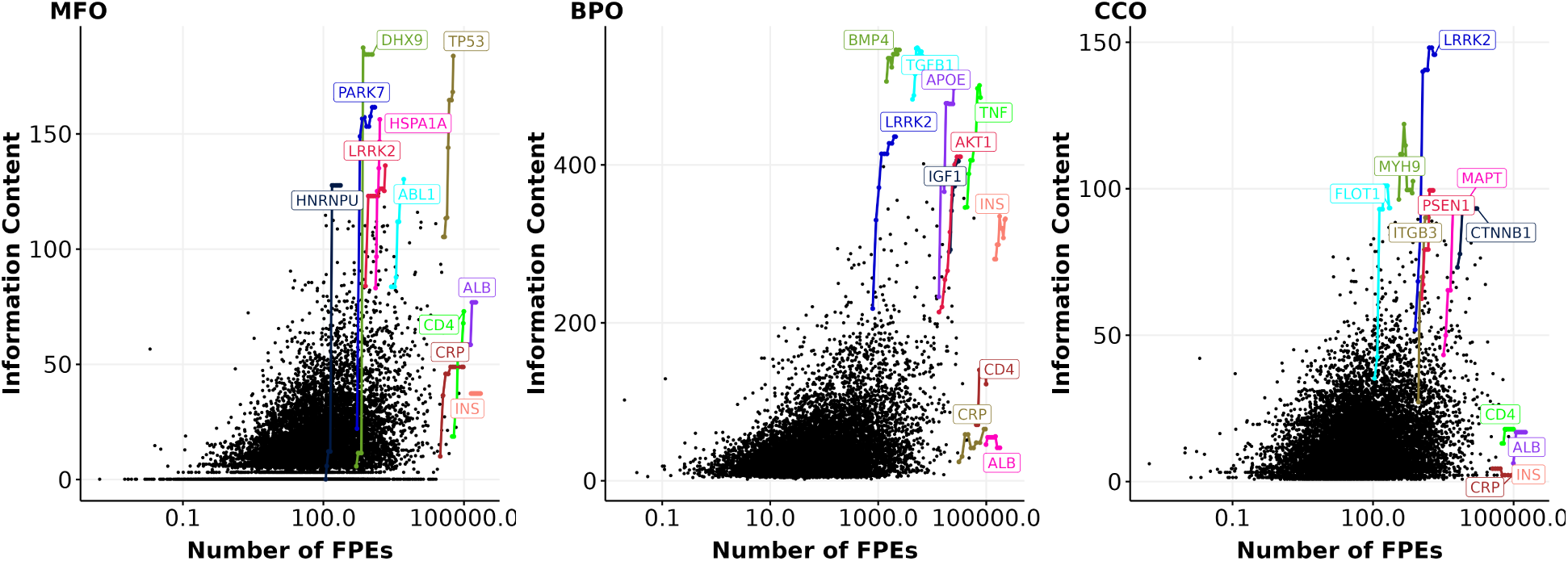
Information content and number of FPEs (in log scale) of proteins in 2022. Each black dot represents the number of FPEs and the information content per protein within an aspect (MFO, BPO, CCO) in 2022. Highlighted genes (colored dots) have one data point for each year from 2013 to 2022. Highlighted lines show the growth in information content and number of FPEs of the most highly annotated proteins and for the four most frequently mentioned proteins from 2013 to 2022 (ALB, CRP, INS, IL6). Gene names encoding for these proteins are shown.

While the correlation between the number of FPEs of human proteins and their information content remains low over time, we discovered that a handful of proteins with certain disease associations displayed a stronger correlation between the number of FPEs and information content. We examined all diseases that had more than 20 associated proteins in an aspect, and performed Spearman correlation tests within each disease. For MFO, this was the case for 61 diseases and for BPO for 54 diseases. After controlling for family-wise error rate of 0.05 using Bonferroni procedure, only four diseases display a significant correlation between the number of FPEs in 2013 and the gain in MFO information content (Figure 5). We did not find any diseases that show a significant correlation between the number of FPEs and the gain in BPO information content of the associated proteins.

**Figure 5:**
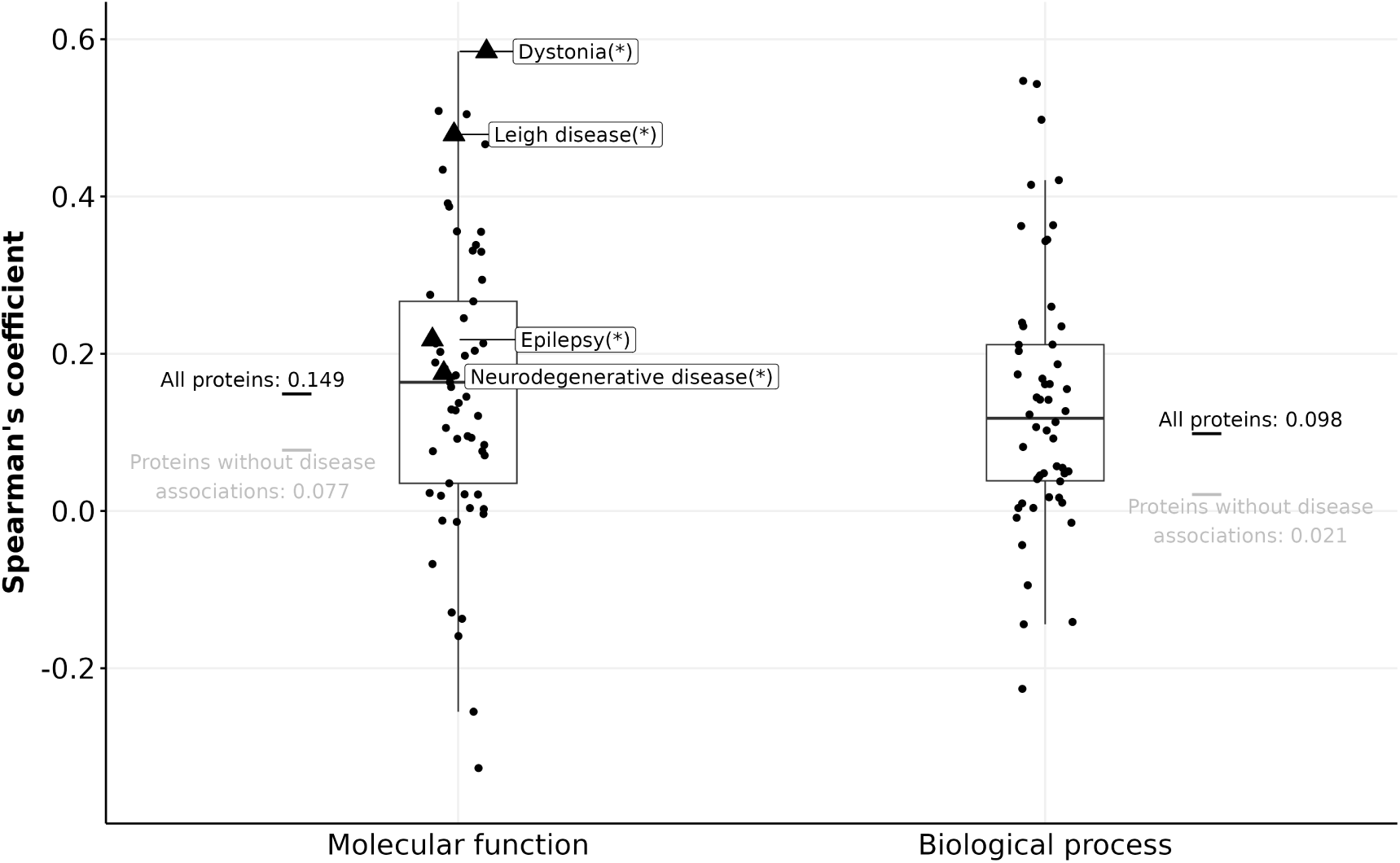
Boxplots showing Spearman’s coefficient between the number of FPEs and the gain in information content (in MFO and BPO) of disease-associated proteins. Triangles (with disease names labeled) indicate diseases with a significant Spearman correlation, and circles indicate diseases without significant Spearman correlation after correcting for multiple testing using the Bonferroni procedure.

For 764 proteins that are associated with dystonia, Leigh disease, epilepsy, or neurodegenerative disease, the correlation between their number of FPEs in 2013 and their gain in MFO information content is stronger than the rest of the proteins (*ρ*_Spearman_ = 0.149). However, given the relatively weak correlations of the four mentioned diseases (*ρ*_Spearman_ *<* 0.6), it is not feasible to predict the gain in information content of protein solely based on its number of FPEs ten years prior. The gain in information content is not predictable using past information content values, nor by using the past number of FPEs.

We hypothesized that the weak correlation between information content of proteins and their number of FPEs could be due to the delay in curation time: the time it takes to compile the information from the research papers into the databases. In particular, biocurators may not review all relevant information from scientific articles for GO annotations because (i) the volume of publications each year is too large, and/or (ii) there are different content priorities for biocuration [41,49,73,74]. We computed the time delay in curation of GO annotations by comparing the annotation timestamp with the publication timestamp of the referenced PubMed article. Figure 6A illustrates that by the end of 2022, the median delay time in experimental annotations in human GO annotation was four years.

**Figure 6:**
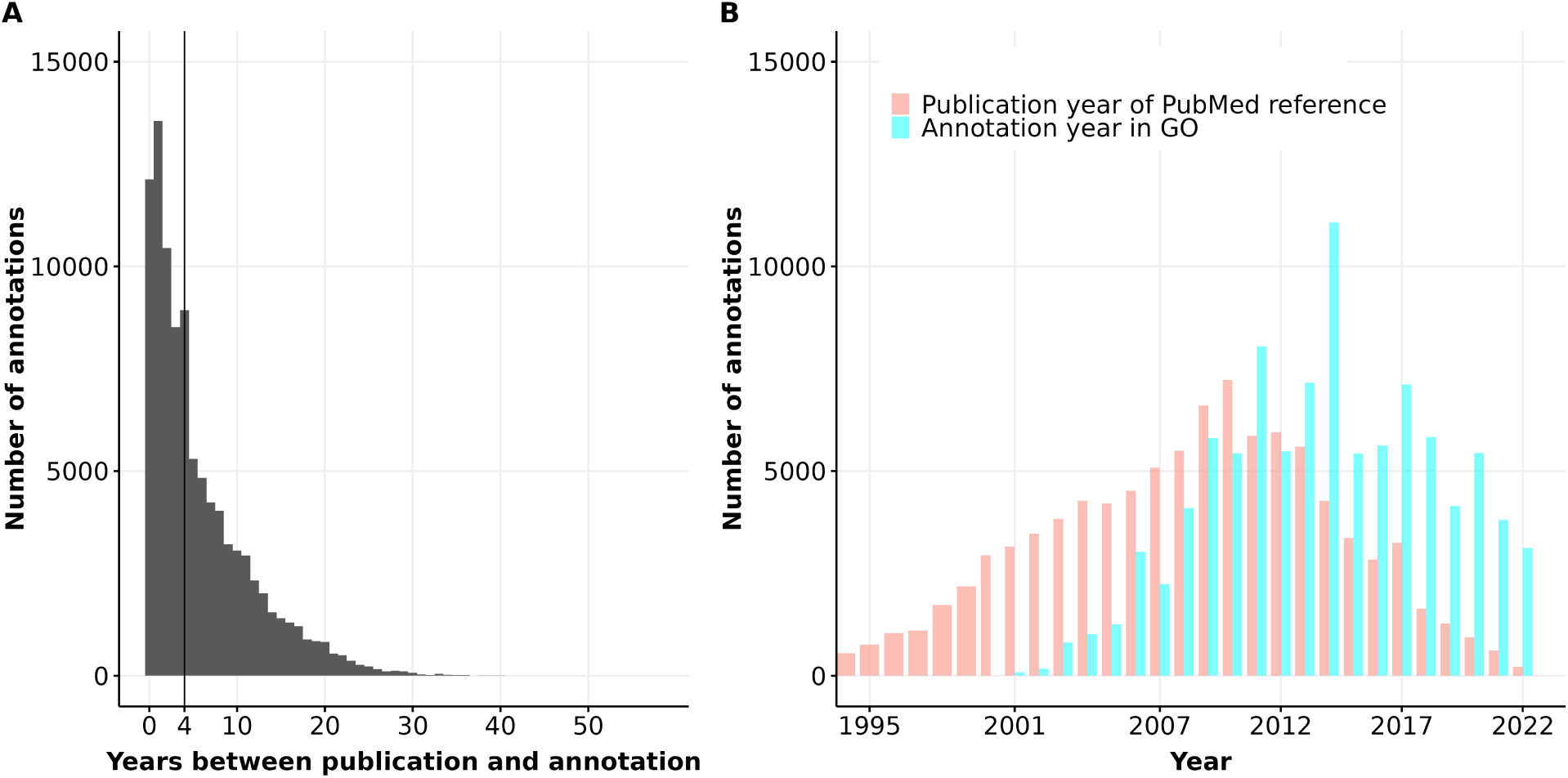
(A) Barplot of the number of years between annotation and publication for experimental GO annotations of human proteins. The vertical line shows the median of this distribution at 4 years. (B) Barplot of the distribution of publication year and annotation year for GO experimental annotations of human proteins.

Figure 6B reveals that the number of experimental annotations curated into human GO annotation files is approximately 4,000 to 7,000 per year from 2008 to 2022, with an exception of nearly 12,000 annotations in 2014. Notably, most experimental annotations for human proteins were derived from articles published between 2008 and 2012. Since then, the number of annotations derived from articles published after 2012 has declined steadily while the number of annotations added to GOA files every year did not decrease. These trends suggest that the pace of curation is not keeping up with the increasing rate of publication, and the median delay in curation is likely to continue to rise as the volume of publications grows.

## 4 Discussion

In this work, we developed metrics to quantify the knowledge about protein function using several GO-based metrics, each of which captures different facets of knowledge about protein function. We used these metrics to quantify the imbalances that exists in our knowledge of the human proteome. Strikingly, we have discovered several types of gaps that persisted over time. Those include an inequality of annotations as found in the GO database, an inequality of interest in different proteins as found in literature mentions, and a time gap between when a protein is mentioned in a publication and when its function is added to the GO knowledgebase.

### 4.1 Accurate quantification of knowledge about protein function is challenging yet crucial

Protein functions are described using GO terms, which are represented in the Gene Ontology with complex biological relationships between different GO terms. Accurately representing and comparing the functions across proteins is challenging due to the intricate nature of GO term relationships, yet it is crucial for studying the biases that exist in the knowledge about protein function. To quantify knowledge about protein function derived from GO annotations, we used term count, unique term count, and information content. The *term count* of a protein is the number of annotations of that protein in a GO annotation file. While term count is useful for capturing additional details about some MFO annotations (e.g., each `protein binding` or `RNA binding` annotation of the same protein specifies a different binding partner), a high value of term count does not imply that the protein performs multiple functions. We thus used *unique term count* to capture the number of unique functions performed by a protein. While unique term count metric reflects the wide variety of functions performed by a protein, it does not capture how informative each function is. *Information content* considers the GO structure and accurately reflects the informativeness of different GO terms. Since the information content of GO terms varies broadly (see Figure S8), unique term count per protein and information content per protein are not equivalent, even though these two metrics are highly correlated (Figure S3) [32].

### 4.2 Information content of certain proteins shows signs of saturation

When quantifying the knowledge we have about protein function over time, we found that some proteins did not gain more information content in recent years. The gain in knowledge, represented by the increase in the information content derived from GO annotations of proteins, is negatively correlated with the initial knowledge in 2013 (*ρ*_Spearman_ = −0.24 and −0.41 in the MFO and BPO aspect, respectively, see Figure S7). One possible explanation for the negative correlation between initial knowledge and the gain in knowledge of proteins is information saturation. We hypothesize that proteins will reach a maximal value of information content in each GO aspect when our knowledge about the protein function is no longer sufficiently captured in GO terms. This is due to the fact that the Gene Ontology only holds partial information about the proteins [75]. It is important to note that information on protein functions is not fully captured in GO annotation files since not all knowledge about protein functions is translated to GO terms [76]. For example, DHX9 has had a burst increase of GO terms since 2017 in both MFO and BPO. However, the information content of DHX9 had plateaued even though we keep gaining new literature-based knowledge about the function of this protein. In particular, the involvement of DHX9 in neurodevelopmental disorder and several hallmarks of cancer are not captured by GO annotations because (1) these roles are due to an interaction of DHX9 with other partners and (2) GO is not a disease ontology [77,78]. Furthermore, the post-transcriptional modification is considered a key step for the protein to obtain its function, but this piece of functional knowledge is out of the scope of GO [79, 80]. Another example is protein Glutaredoxin-3 (GLRX3). As new knowledge about this protein was discovered, its annotations in MFO remained the same since 2017, and thus, its information content did not change because the Gene Ontology does not have a term to capture more detailed findings on its molecular function. This is the case with most proteins where the knowledge about functions is not captured fully by experimental GO annotations [20, 51]. Therefore, the total knowledge about protein functions is not reflected through Gene Ontology annotations alone, which gives rise to the aforementioned plateauing of information content measured from GO annotations of the proteins. Nevertheless, GO annotations remain useful for large-scale computational studies due to their widespread use and computational amenability. The partial knowledge represented in GO highlights the need for more comprehensive quantification methods across several knowledgebases and ontologies to better capture knowledge about protein functions in computational analyses [76].

Although the Gene Ontology provides partial knowledge about protein function and we may eventually exhaust GO terms to annotate certain proteins, it remains challenging to test the hypothesis that all proteins will reach saturation in GO information content. As new technologies emerge, we continue to discover additional GO terms and update the GO structure, enabling new annotations of proteins that previously appeared to reach saturation [22, 52, 81]. While the rate of new GO terms being added has slowed down in recent versions of the GO, it is almost guaranteed that the current Gene Ontology will not remain static [41].

### 4.3 Annotation and publication inequalities remain consistently high

Following the quantification of “knowledge wealth” of human proteins using GO-based metrics, we examined the gap that existed in the knowledge about protein function among proteins in the human proteome. A previous study by Haynes *et al* reported that Gini coefficients of protein function annotations increase over time in which the annotation inequality in GO (as computed from unique term count) rises from 0.25 in 2001 to 0.47 in 2017 [23]. Specifically, Figure S1D in the same study has shown that the Gini coefficients fluctuate around 0.5 from 2013 to 2017 for experimental annotations, which is the time period that overlaps with our analysis. We also found that Gini coefficients of unique term count of proteins fluctuate from 2013 to 2017, with values of approximately 0.4, 0.5, and 0.4 in the aspects of MFO, BPO, and CCO, respectively (Figure 2). The slight disparity in Gini coefficients between ours and Haynes’ results could be because we distinguished between three GO aspects when calculating Gini coefficients.

Additionally, our investigation of earlier-annotated proteins (annotated in or before 2013) reveals that the MFO information content distribution is less unequal from 2013 to 2022, as reflected in the decreasing trend of the Gini coefficient in this period (Figure 2B). In fact, nearly half of the proteins that were only annotated with `protein binding` (or “protein-binding-only“ proteins) have gained more informative annotations since then. The gain in information content from the protein-binding-only subset of proteins is influential to the inequality of information content of proteins annotated in or before 2013. In other words, the inequality for the set of earlier-annotated proteins was greatly decreased by discovering more informative molecular functions for lesser annotated proteins. This observation strongly suggests that focused research could expand the knowledge of less annotated proteins, particularly in the protein-binding-only subset. Less annotation effort being spent on well-known proteins implies that more annotation effort will go to lesser-known earlier-annotated proteins. Therefore, it is crucial to implement comprehensive annotation efforts for less-annotated proteins. Focused research and increased resources should be directed towards these under-annotated proteins, especially proteins that were under-annotated for several years, to reduce overall annotation inequality and ultimately advance our knowledge of human proteomics.

On the other hand, the decreasing trend in Gini coefficient was not seen in BPO information content and CCO information content (Figure S4), as there is no low-information term similar to `protein binding` in these two aspects that can be improved to more informative GO annotations for proteins. Therefore, it is harder to decrease the inequalities in information content in the aspects of BPO and CCO.

We were also interested in the gap in “interest wealth”, the gap that exists in the attention given to different proteins in the literature. To do so, we calculated Gini coefficients using the number of FPEs, which is the interest metric in this study. We observed a high inequality in literature mentions of human proteins, with a Gini coefficient of 0.87 decreasing to 0.85 over the decade, indicating a strong bias in research efforts towards a small number of proteins. This finding aligns with previous results [19–21]). However, as we see, the inequality in knowledge about protein function, ranging from 0.4 to 0.65 (as captured by the GO-based metrics), is not nearly as high as the inequality in interest. In other words, although research efforts are heavily biased towards more popular proteins, GO annotations of the proteins seem to be assigned more equally. This could be explained by three reasons: (i) biocurators assign annotations to proteins more comprehensively, resulting in higher and more equal coverage of the human proteome compared to the study of these proteins [82], (ii) the knowledge of protein function captured in GO is limited, leading some proteins to reach saturation in GO-based knowledge while still accruing interest through publications, or (iii) part of the disparity in inequalities may be due to a larger underlying protein population used in interest analyses compared to knowledge analyses (i.e., the number of proteins mentioned in at least one literature is greater than the number of proteins with at least one experimental annotation).

### 4.4 The gap between knowledge of and interest in human proteins impacts ongoing research

We hypothesized that there should be a correlation between the gain in knowledge and the initial interest because proteins with more mentions 10 years ago, accrued more mentions and more annotations in the future. However, we found that the gain in knowledge is not predicted by the initial interest in 2013 (*ρ*_Spearman_ = 0.11 for gain in MFO information content and *ρ*_Spearman_ = 0.05 for gain in BPO information content). The weak correlation between the interest in proteins in 2013 and the gain in knowledge of protein function after ten years is surprising since we expected a protein that is studied more should also acquire more annotations. The reason for the weak correlation between the initial interest (number of FPEs in 2013) and the gain information content is that the number of FPEs of these proteins also includes publications that do not provide function annotation. In other words, the number of FPEs reflects the total knowledge about proteins, not limited to protein function annotations. Poux *et al* have shown that only 35%-45% of PubMed is relevant for UniProtKB/Swiss-Prot curation, i.e., these publications contain protein sequence and functional information [49]. GO annotations of protein functions occupy an even lower percentage of all publications about proteins since redundant publications about the same function of a certain protein do not provide new GO terms for that protein. In addition, when a publication discovers a new function for a protein, the evidence is not always sufficient to curate this information into the database [49]. Therefore, the number of publications that provide GO annotations for proteins, which gives rise to the information content of proteins, is only a small fraction of the publications which we derived the number of FPEs from. Additionally, we characterized the time gap between publication and database curation, contributing to the disparity between research interest and database-store knowledge. We showed that the experimental annotations, which are of high quality and time-consuming to curate, have a median delay of four years from the point of publication to annotation, with 25% delayed by ten years or more (Figure 6). For proteins that experience a delay greater than ten years, significant functional discoveries in literature ten years ago may be absent from current GO annotations, explaining part of the gap between the interest in these proteins and the knowledge of their function. This delay in biocuration in which dozens of current experiments not being curated into GO knowledgebase yet could hinder ongoing research, especially for lesser known proteins [15, 83, 84]. These findings imply that researchers may be making sub-optimal decisions when prioritizing which proteins to study based on publication history, as the publication gain does not aid the discovery of new knowledge effectively, or the knowledge about protein functions was not adequately captured in their GO annotations. In essence, although the initial interest predicts the gain in interest very well, it is not a strong indicator for knowledge gain. Therefore, more comprehensive approaches are needed to capture other aspects of knowledge on protein functions and the attention these proteins receive in the field. The question that remains is what factors determine the gain in information content for proteins.

## 5 Conclusions

Here we provided a survey of the knowledge about, the interest in, and the correlation between knowledge of and interest in the human proteome between 2013-2022. We examined the changes in inequality in the knowledge and interest distribution among human proteins over this 10-year period. We have shown that the inequality in the knowledge about protein function, measured by information content, is consistently high, which reflects a bias towards a few “popular” proteins. However, these highly annotated proteins gain less information content over time, with several proteins showing potential signs of information saturation. By directing more annotation efforts to the proteins that carry little information, the high inequality in knowledge about protein function can be mitigated. Additionally, we show that the research interest in proteins, quantified by the number of normalized mentions in publications, also has a higher inequality than the inequality measured using the GO annotation of protein function. Furthermore, we noted that the median time delay between the publication about protein function and the curation of experimental GO annotations is four years, highlighting a gap between the information available in scientific publications and the information captured in curated databases such as GO. It is imperative we address these delays and biases, and develop a productive system for protein function annotation. We suggest developing more systematic approaches to assess the total knowledge about protein functions, and a refocusing of annotation efforts on less studied proteins.

## Code and Data Availability

The code used to extract data from GOA files and subsequent analyses in this study is available on GitHub at https://github.com/FriedbergLab/protein-annotation-bias.

Processed datasets (knowledge and interest metrics of proteins from 2013 to 2022) are available in the Figshare folder at https://doi.org/10.6084/m9.figshare.27252825. External data files required in the analysis are available at https://doi.org/10.6084/m9.figshare.27252282.

## Acknowledgements

To be entered after acceptance.

