## Supplemental Materials for "A Longitudinal Analysis of Function Annotations of the Human Proteome Reveals Consistently High Biases"

### Supplementary Materials for A Longitudinal Analysis of Function Annotations of the Human Proteome Reveals Consistently High Biases

An Phan, Parnal Joshi, Claus Kadelka, Iddo Friedberg

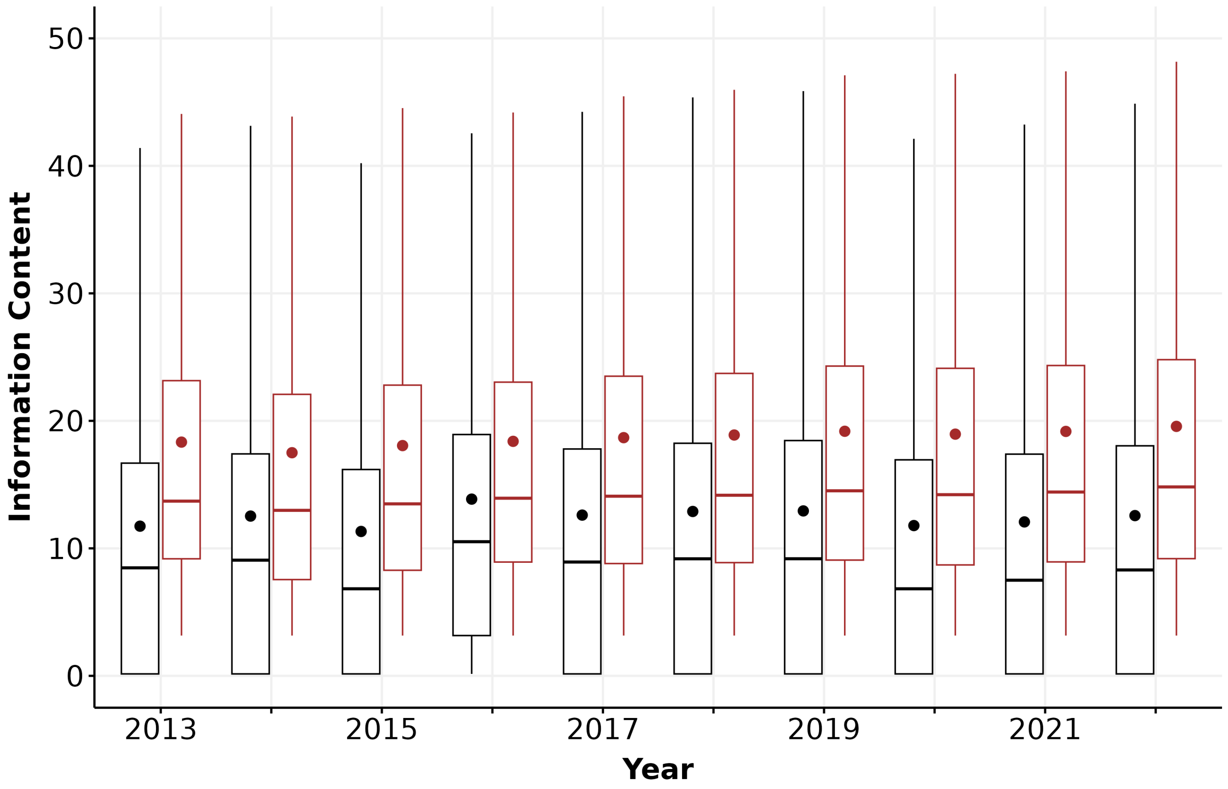


**Figure S1: Boxplot showing distribution of information content of all proteins (in black) and the set of proteins excluding protein-binding-only proteins (in brown) in the Molecular function aspect.** Each box extends across the interquartile range (IQR), vertical lines extend to the lowest data point (or highest data point) still within 1.5 IQR of the lower quartile (or the upper quartile), a horizontal line shows the median and a dot shows the mean value. Outliers are not displayed.

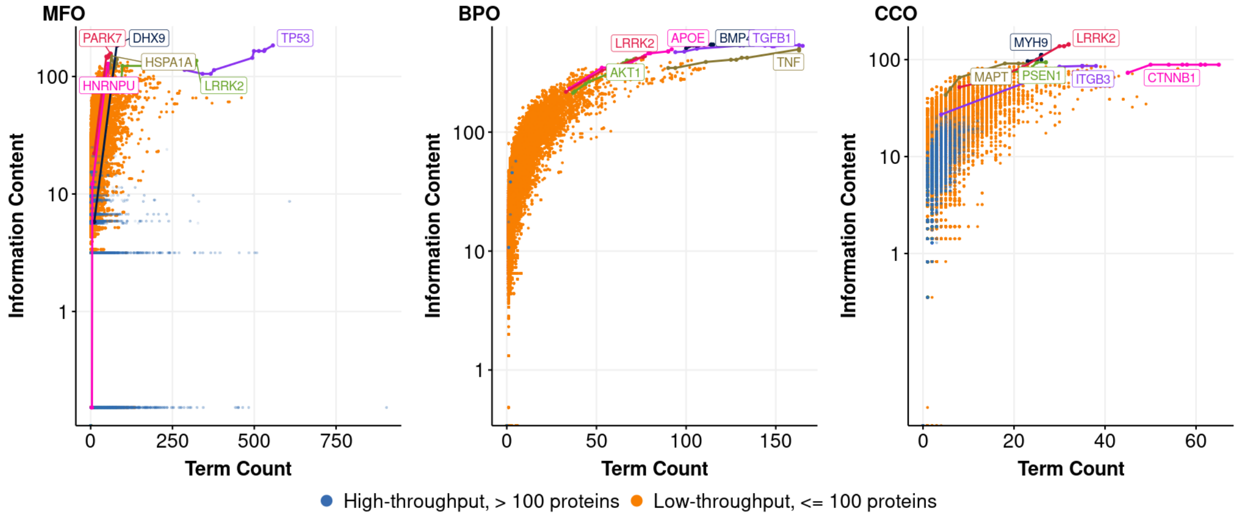


**Figure S2: Information content (in log scale) and term count of proteins studied experimentally from 2013 to 2022.** Each dot represents the term count and the information content per protein at a time point in Molecular Function (MFO), Biological Process (BPO), and Cellular Component (CCO). Highlighted lines show the growth in information content and unique term count of the most highly annotated proteins over time in low-throughput experiments (gene names encoding for these proteins are labeled).

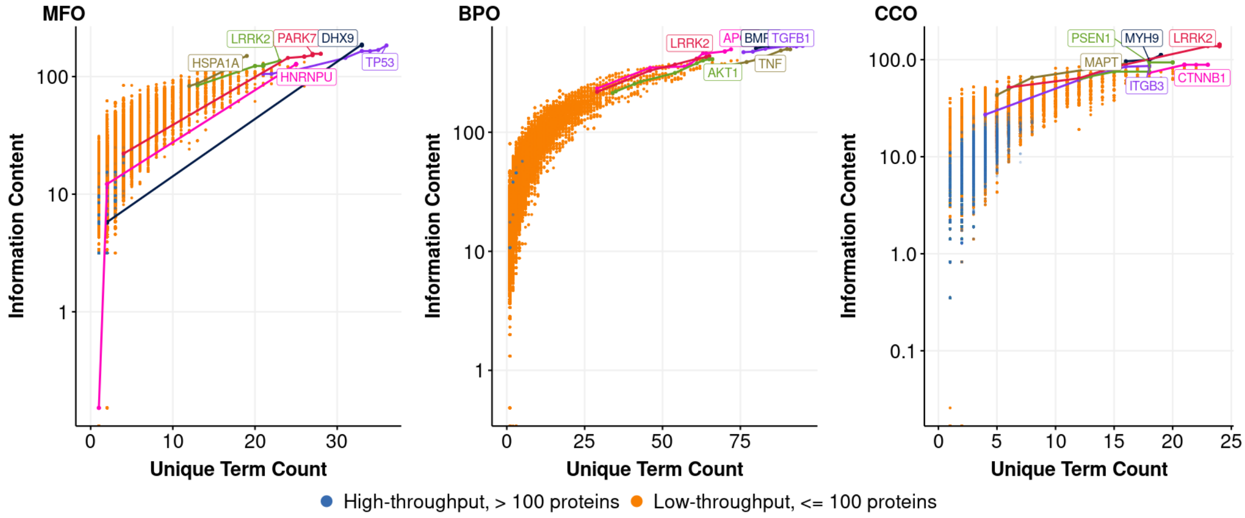


**Figure S3: Information content (in log scale) and unique term count of proteins studied experimentally from 2013 to 2022.** Each dot represents the term count and the information content per protein at a time point in Molecular Function (MFO), Biological Process (BPO), and Cellular Component (CCO). Highlighted lines show the growth in information content and unique term count of the most highly annotated proteins over time in low-throughput experiments (gene names encoding for these proteins are labeled).

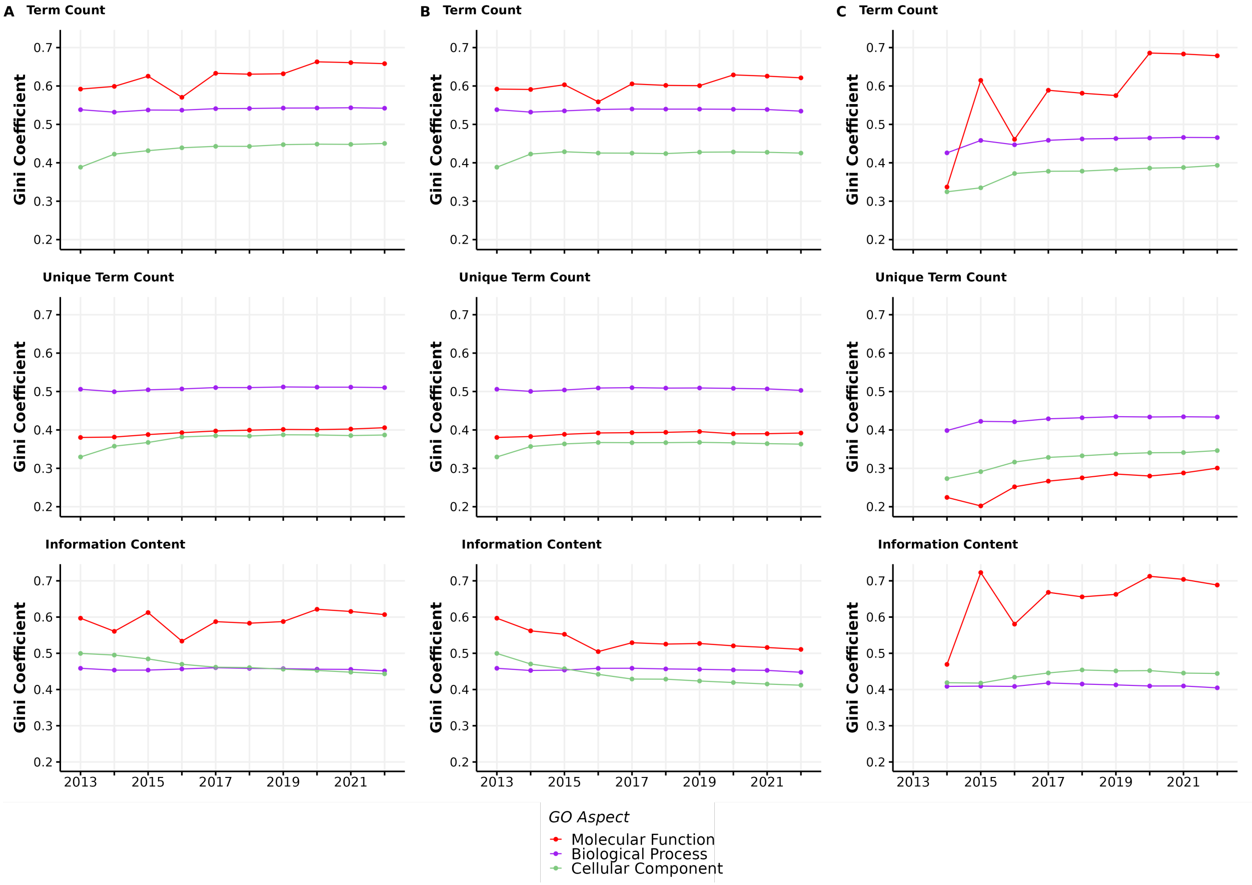


**Figure S4: Gini coefficients (0-1) from 2013-2022 calculated from three knowledge metrics (term count, unique term count, and information content) across three GO aspects: Molecular function (MFO, in red), Biological process (BPO, in purple), and Cellular component (CCO, in green).** Panel A includes all proteins experimentally annotated in each aspect. Panel B includes proteins annotated in or before 2013, and panel C only includes proteins annotated after 2013.

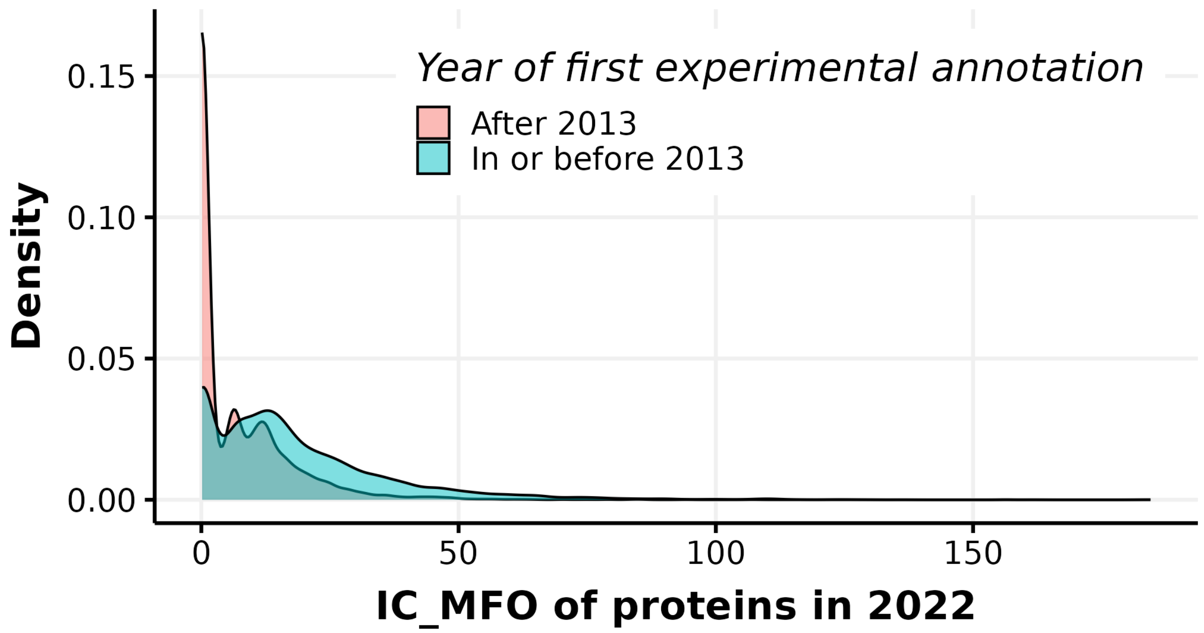


**Figure S5:** **Density plots showing information content in MFO (IC_MFO) in 2022 of two subsets of proteins: first experimentally annotated in or before 2013, and first experimentally annotated after 2013.**

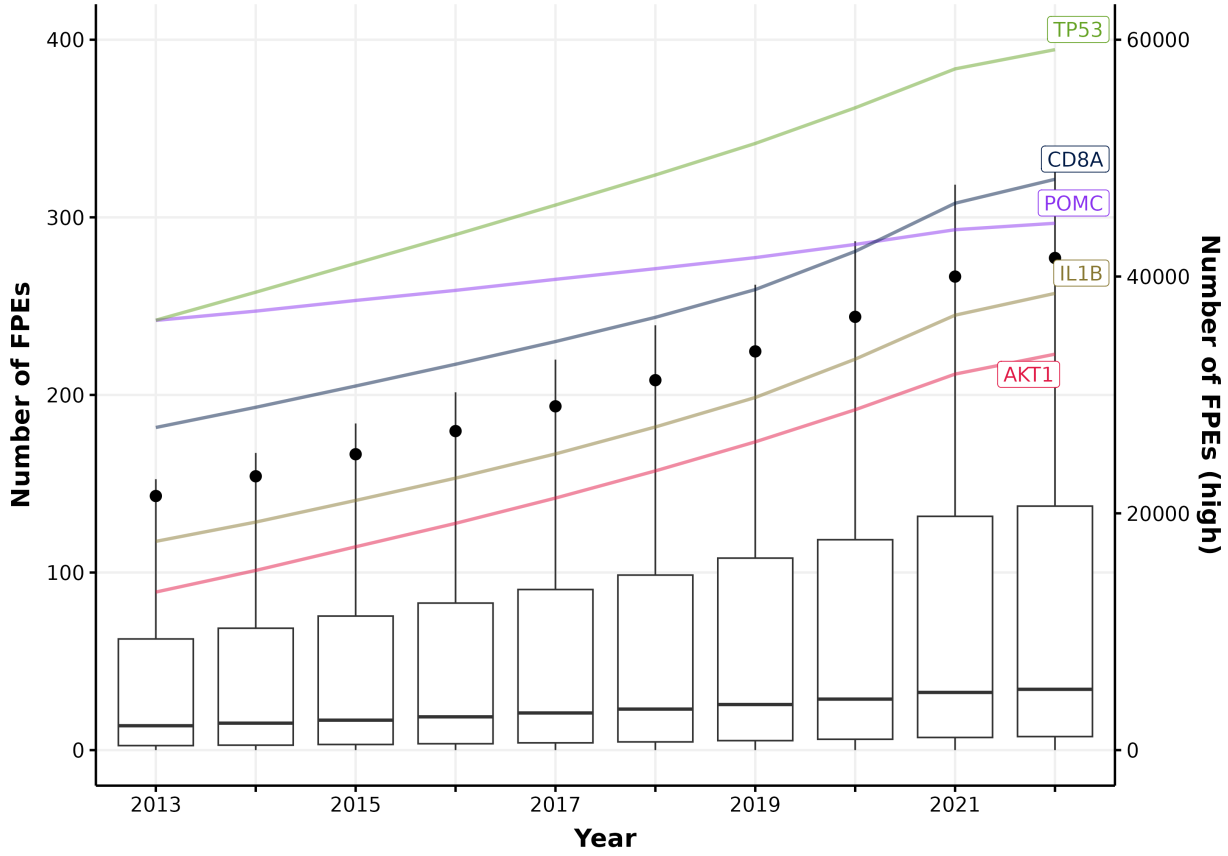


**Figure S6: Boxplots showing number of FPEs of proteins from 2013 to 2022.** Circles indicate the average number of FPEs of all proteins for each year. Highlighted lines show proteins that have the highest cumulative number of FPEs up to 2022 whose values are shown in the right y-axis of "Number of FPEs (high)". Proteins that are mainly used as clinical assays have been removed.

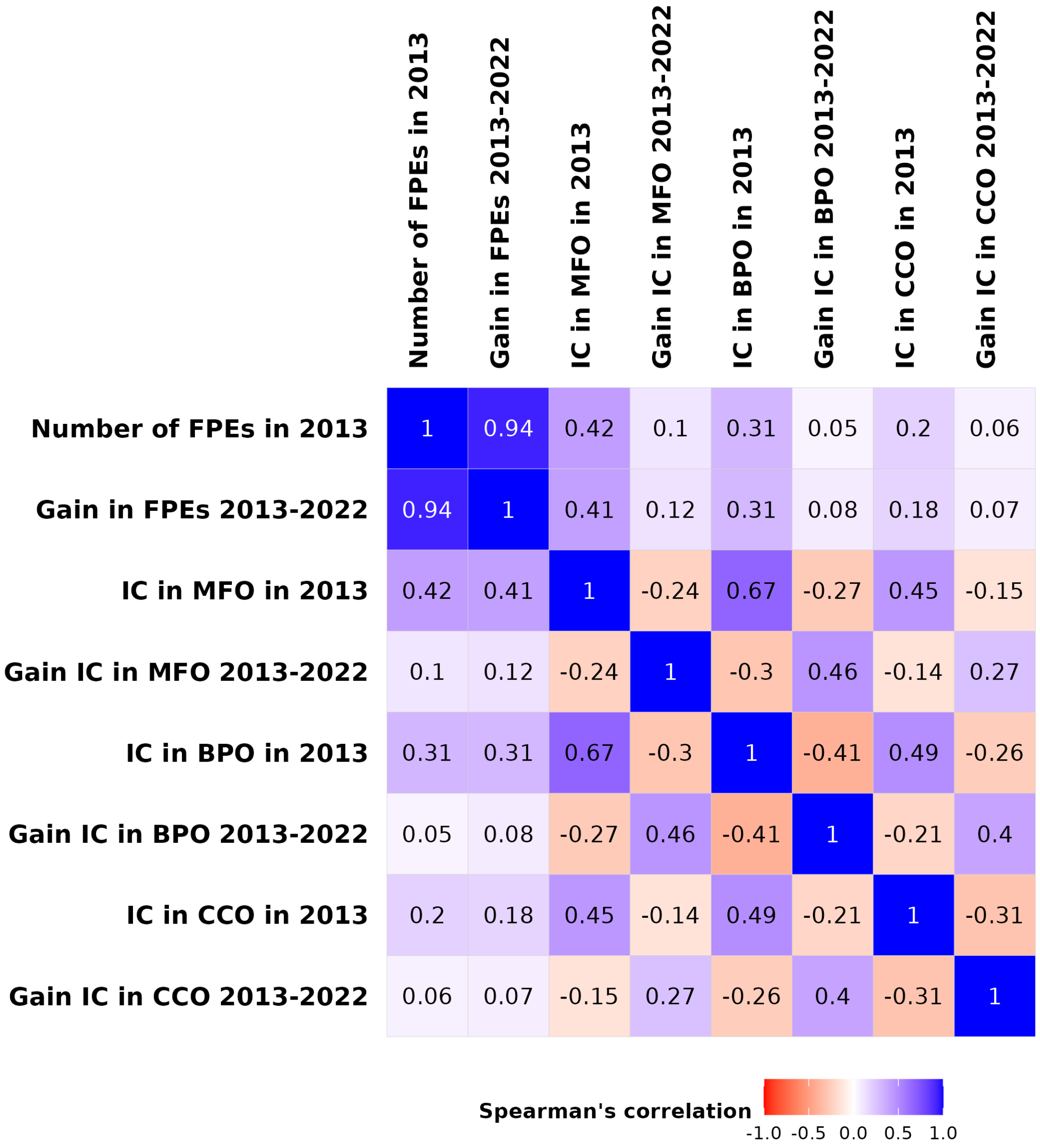


**Figure S7:** **Heatmap showing Spearman correlation between number of FPEs, information content in every aspect in 2013, and the gain of these metrics from 2013 to 2022.**

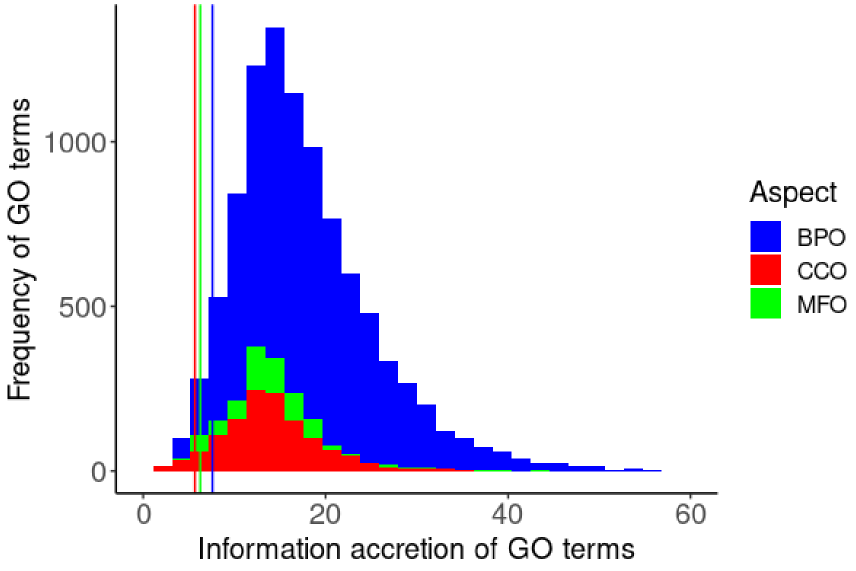


**Figure S8: Histogram of information content of GO terms in three aspects of the GO.** The vertical lines indicate the 5th percentile, approximately at 6 bits, of the distribution in corresponding color. MFO: Molecular function, BPO: Biological process, CCO: Cellular component.

**Table S1. Number of proteins and the median term count and information content by year in all GO aspects. MFO: Molecular function, BPO: Biological process, CCO: Cellular component.**

| | Number of proteins | | | Unique term count | | | Information content | | |
|---|---|---|---|---|---|---|---|---|---|
| Year | MFO | BPO | CCO | MFO | BPO | CCO | MFO | BPO | CCO |
| 2013 | 7831 | 5856 | 9023 | 3 | 2 | 2 | 8.44 | 24.48 | 5.74 |
| 2014 | 8953 | 6887 | 11348 | 3 | 3 | 2 | 9.07 | 25.23 | 5.74 |
| 2015 | 10946 | 7598 | 12469 | 4 | 3 | 2 | 6.74 | 26.59 | 5.96 |
| 2016 | 9414 | 7619 | 10056 | 3 | 3 | 2 | 10.51 | 26.36 | 7.94 |
| 2017 | 11747 | 8393 | 10865 | 4 | 3 | 3 | 8.88 | 27.01 | 8.6 |
| 2018 | 12000 | 8651 | 10913 | 4 | 3 | 3 | 9.17 | 27.43 | 8.94 |
| 2019 | 12253 | 9041 | 11193 | 4 | 3 | 3 | 9.18 | 27.8 | 9.16 |
| 2020 | 14238 | 9304 | 11340 | 6 | 3 | 3 | 6.83 | 28.13 | 9.45 |
| 2021 | 14335 | 9436 | 11466 | 6 | 3 | 3 | 7.5 | 28.45 | 9.5 |
| 2022 | 14544 | 9869 | 11791 | 7 | 3 | 3 | 8.3 | 29.25 | 10.27 |

**Table S2. List of proteins with highest information content in at least one GO aspect with UniProtKB identifier, gene name, and protein name.**

| UniProtKB Identifier | Gene Name | Protein Name |
|---|---|---|
| Q08211 | DHX9 | ATP-dependent RNA helicase A |
| P02649 | APOE | Apolipoprotein E |
| P12644 | BMP4 | Bone morphogenetic protein 4 |
| P35222 | CTNNB1 | Catenin beta-1 |
| P04637 | TP53 | Cellular tumor antigen p53 |
| P0DMV8 | HSPA1A | Heat shock 70 kDa protein 1A |
| Q00839 | HNRNPU | Heterogeneous nuclear ribonucleoprotein U |
| P05106 | ITGB3 | Integrin beta-3 |
| Q5S007 | LRRK2 | Leucine-rich repeat serine/threonine-protein kinase 2 |
| P10636 | MAPT | Microtubule-associated protein tau |
| P35579 | MYH9 | Myosin-9 |
| Q99497 | PARK7 | Parkinson disease protein 7 |
| P49768 | PSEN1 | Presenilin-1 |
| P31749 | AKT1 | RAC-alpha serine/threonine-protein kinase |
| P01137 | TGFB1 | Transforming growth factor beta-1 proprotein |
| P01375 | TNF | Tumor necrosis factor |
